# Stable host gene expression in the gut of adult *Drosophila melanogaster* with different bacterial mono-associations

**DOI:** 10.1101/053512

**Authors:** Carolyn Elya, Vivian Zhang, Will Ludington, Michael B Eisen

## Abstract

There is growing evidence that the microbes found in the digestive tracts of animals influence host biology, but we still do not understand how this comes about. Here, we evaluated how different microbial species commonly associated with laboratory-reared *Drosophila melanogaster* impact host biology at the level of gene expression in the dissected adult gut or the entire adult organism. We observed that guts from gnotobiotic animals associated from the embryonic stage with either zero, one or three bacterial species demonstrated indistinguishable transcriptional profiles. Additionally, we found that the gut transcriptional profiles of animals reared in the presence of the yeast *Saccharomyces cerevisiae* alone or in combination with bacteria could recapitulate those of conventionally-reared animals. In contrast, we found whole body transcriptional profiles of conventionally-reared animals were distinct from all of the gnotobiotic treatments tested. Our data suggest that adult flies are insensitive to the ingestion of different bacterial species but that prior to adulthood, different microbes impact the host in ways that lead to global transcriptional differences observable across the whole adult body.

## Introduction

The digestive tracts of virtually all animals examined to date are inhabited by microbes and there is increasing evidence that interactions between gut microbes and their animal hosts influence a wide range of host phenotypes [1]. However the molecular basis for these effects is poorly understood.

*Drosophila melanogaster* has become a model for the study of host-microbe interactions in the gut owing to the relative simplicity of the tissue and its microbial community. The *Drosophila* gut microbe community is comprised of about 50 bacterial species in wild flies [2–5] and about ten species in lab-reared flies [2,3,6–8]. In lab-reared animals, the majority of these bacteria are lactic acid and acetic acid producers (members of family Lactobacillaceae and Acetobacteraceae, respectively) that can be cultured outside of the host. *Drosophila* embryos can be stripped of their endogenous microflora using household bleach [9] and either maintained under sterile conditions to generate axenic (germ-free) animals or treated with a defined set of microbes to generate gnotobiotic (of known microbial content) animals [10].

A variety of phenotypes have been associated with the presence and/or specific composition of microbes in the *Drosophila melanogaster* gut including nutrition and metabolism [8,10,11], intestinal cell growth [12–14], development [15,16], lifespan [7,17–19] and a variety of different behaviors, including social attraction in larvae and assortative mating in adults of two lab-reared strains [20–22].

The presence of microbes in the environment and gut has a significant effect on physiology, morphology and gene expression in the adult *D. melanogaster* gut [11,19,23], however, the effect of individual microbe species on the gene expression of adult guts has not been examined. Given the known differential phenotypic effects of different bacterial taxa, we hypothesized that the epithelial cells of the digestive tract might respond to different bacteria in different ways, and that these responses might provide clues to the molecular mechanisms that underlie the effects of specific microbes of *Drosophila* physiology and behavior. Here we describe the results of a series of experiments using mRNA sequencing to analyze gene expression in the guts of a laboratory line of *D. melanogaster* reared from embryogenesis to early adulthood with different combinations of bacteria and yeast commonly found in lab populations.

## Results

### Limited variation in transcription in adult *D. melanogaster* gut in response to mono-association with different bacterial species

To elucidate the effects of specific bacterial species on host gut gene expression we implemented previously published protocols [10] to mono-associate laboratory stocks of *D. melanogaster* with three bacterial species previously shown to associate with *D. melanogaster* in the lab: *Acetobacter pasteurianus, Lactobacillus brevis* and *Lactobacillus plantarum. A. pasteurianus* and *L. brevis* were selected based on their high abundance in a survey of wild-type female (CantonS) flies reared on standard media in our lab (Figure S1) as well as their ubiquity in other gut microbe surveys of *D. melanogaster* [2,3,6–8]. *L. plantarum* was chosen because it has been observed in several other gut microbe surveys and has been implicated in mediating specific aspects of fly development and behavior [16,22].

Briefly, sterile embryos of *D. melanogaster* (CantonS) were placed in vials inoculated with 5x106 CFUs of the relevant bacteria and allowed to develop under standard conditions. At five days after eclosion, gut samples from single female adults were prepared for mRNA sequencing and sequenced using standard methods.

After aligning sequencing reads to the *D. melanogaster* genome and estimating transcript abundance (FPKM) for each transcript, we visually examined variability in gene expression within and between experiments and treatments across samples organized by either experimental date (Figure 1A) or bacterial mono-association (Figure 1B). Contrary to our expectation, we did not observe any significant differences in gene expression associated with the different bacterial treatments.

**Figure 1).**
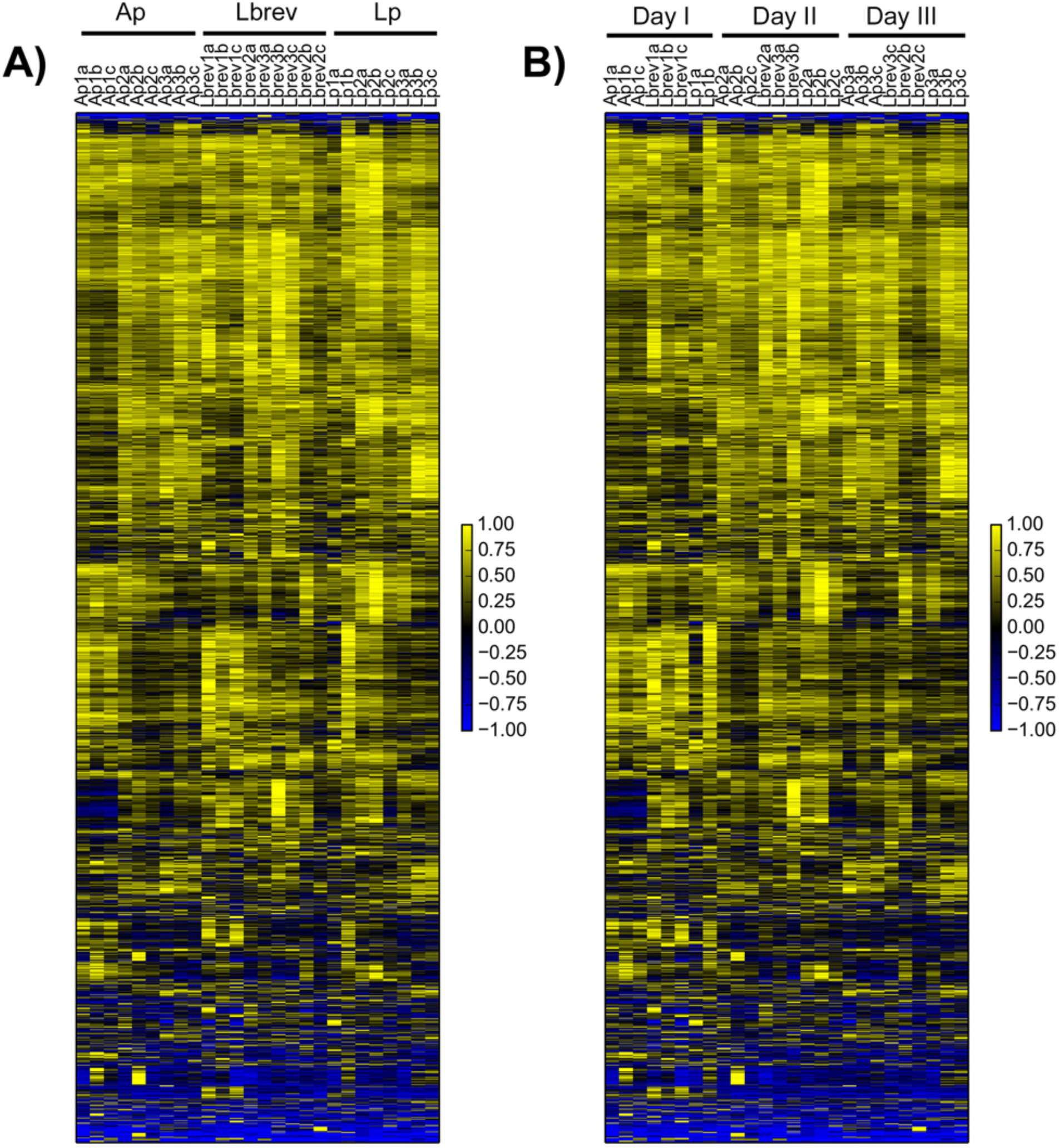
Limited variation in gut gene expression with bacterial mono-association. Expression data from guts dissected from five-day post-eclosion, ***Wolbachia***-free, mated female CantonS ***D. melanogaster*** individuals mono-associated with one of three bacteria (Ap = ***A. pastuerianus***, Lbrev = ***L. brevis***, Lp = ***L. plantarum***) were clustered by gene (average linkage, uncentered correlation) after first filtering out genes that lacked three instances of FPKM greater than two (Gene Cluster 3.0). FPKM values for each gene were normalized to range from-1 to 1 before plotting. A) Samples arranged by bacterial treatment. B) Samples arranged by date of experiment. Scale bars for each heatmap are given to the right of the plot.

To confirm this visual impression, we used a simple statistical test (ANOVA) to identify individual genes differentially expressed between treatments. After applying a Bonferroni correction for multiple testing, the expression levels of only two genes were significantly associated with treatment: CAH2, a carbonic anhydrase, and CG17574, a gene of unknown function (Figure 2). Even relaxing our significance threshold, and considering the fifty genes that demonstrate the most significant differences in expression within this set, it is clear that there are minimal differences in gene expression between the three different bacterial mono-associations (Figure 2). Instead, it appears that each bacterial species alone affected host gene expression in the same way.

**Figure 2).**
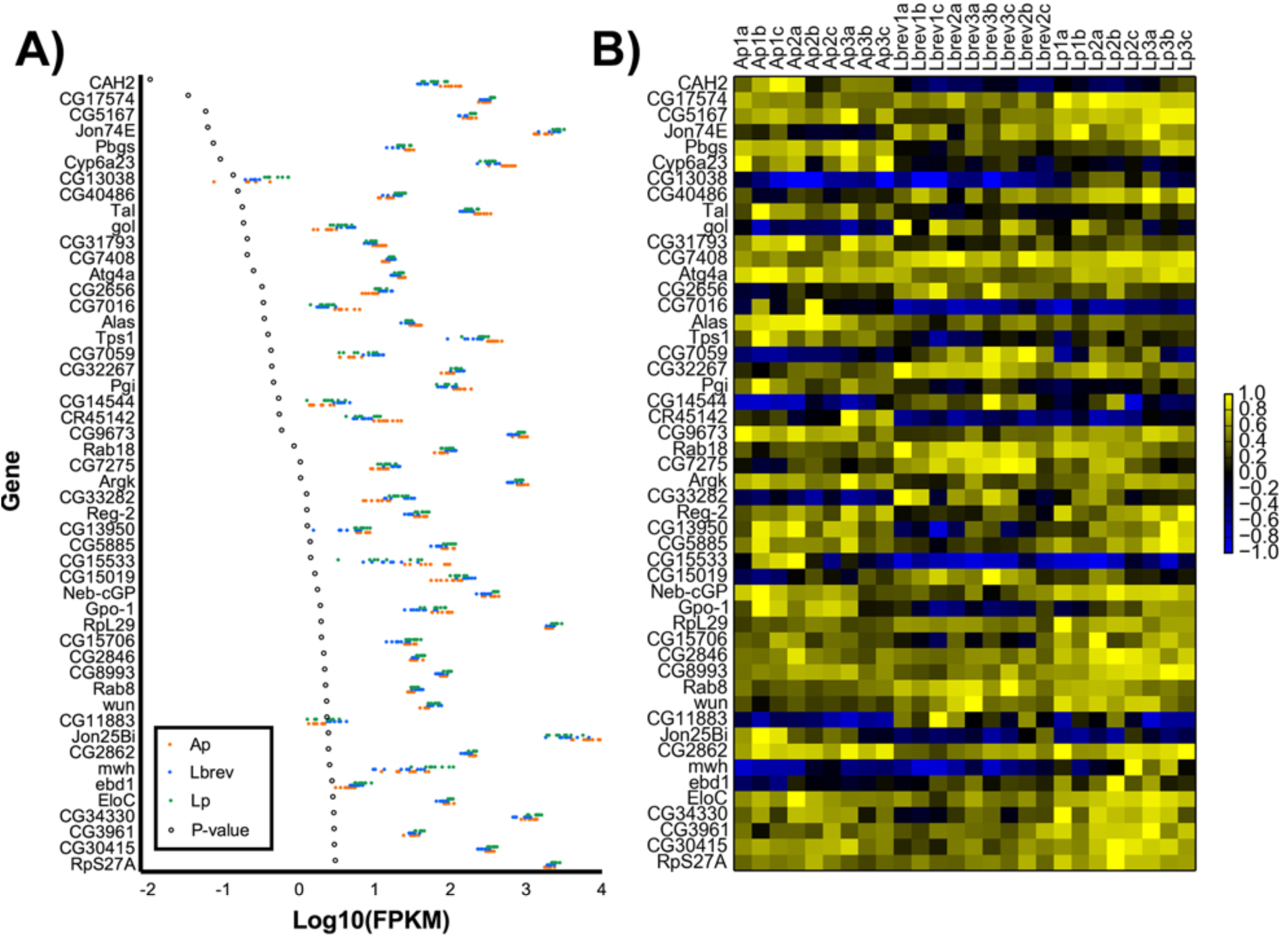
Genes showing greatest difference in expression values from dissected adult guts as determined by one-way ANOVA. A) Scatterplot of log10-transformed FPKM values for each bacteria mono-associated gut replicate (Ap = ***A. pastuerianus***, Lbrev = ***L. brevis***, Lp = ***L. plantarum***). Genes are ordered from lowest ANOVA p-value (top) to highest (bottom). P-values have undergone a Bonferroni correction for multiple testing. B) Data from A presented as a heatmap. FPKM values for each gene are linearly normalized to range from-1 to 1 before plotting.

### Host transcription in the gut is markedly different between conventional and yeast mono-association compared to axenic and bacteria mono-association treatments

After observing a lack of differences in our initial set of samples, we decided to expand our dataset to include other treatments that might shed light on this unexpected result. To this end, we measured gene expression in guts from conventionally reared flies, axenic flies and flies mono-associated with the yeast *Saccharomyces cerevisiae* (which is known to be associated with flies in the wild [24] and can alone provide complete nutrition for the developing fly [25]) rather than bacteria. Additionally, we tested if providing a simplified multi-species microbial community (either all three of *A. pasteurianus, L.brevis* and *L. plantarum* with or without *S. cerevisiae*) would yield a transcriptional program more similar to that of conventional than to mono-associated samples.

Based on studies demonstrating that microbes play critical roles in animal development, we anticipated that the expression pattern of guts from axenic flies would be distinct from those of mono-associated, poly-associated (bearing a simplified microbial community) or conventional animals [11,23]. We also expected expression in mono-associated and poly-associated flies to either closely resemble that of conventionally-reared flies or lie somewhere in between the axenic and conventional samples. Finally, we did not expect that providing yeast, either alone or with all three of our selected bacteria, would have a significant impact on gut gene expression, as previous studies suggest that most vegetative *S. cerevisiae* ingested by *D. melanogaster* do not survive passage through the digestive tract [26,27]. If *S. cerevisiae* is only metabolized and does not actively survive the GI tract, we predicted that providing live *S. cerevisiae* would have no significant impact on the flies, since they already were provided dead *S. cerevisiae* in their diet.

To test these predictions, we dissected and sequenced mRNA from wild-type, 5-day-old, female CantonS flies that were either axenic, conventional, mono-associated with *S. cerevisiae*, poly-associated with an equal amounts of *A. pasteurianus, L. brevis* and *L. plantarum* or poly-associated with an equal amounts of *A. pasteurianus, L. brevis, L. plantarum* with or without *S. cerevisiae*.

As above, we first examined the aggregate gene expression data visually (Figure 3A). In contrast to our expectations, the expression data clearly showed that guts from gnotobiotic animals whose treatment included yeast gave transcription patterns most similar to those of guts from conventional animals. We were further surprised to observe that rather than being distinct from all other samples as we had expected, transcriptional patterns of guts from axenic animals most closely resembled that of animals either mono-or poly-associated with bacteria. Taken together, samples appeared to fall within one of two transcriptional regimes: “conventional-like” ( conventional and yeast-mono and poly-associations) and “bacteria-like” (axenic, bacteria mono-and poly-associations).

**Figure 3).**
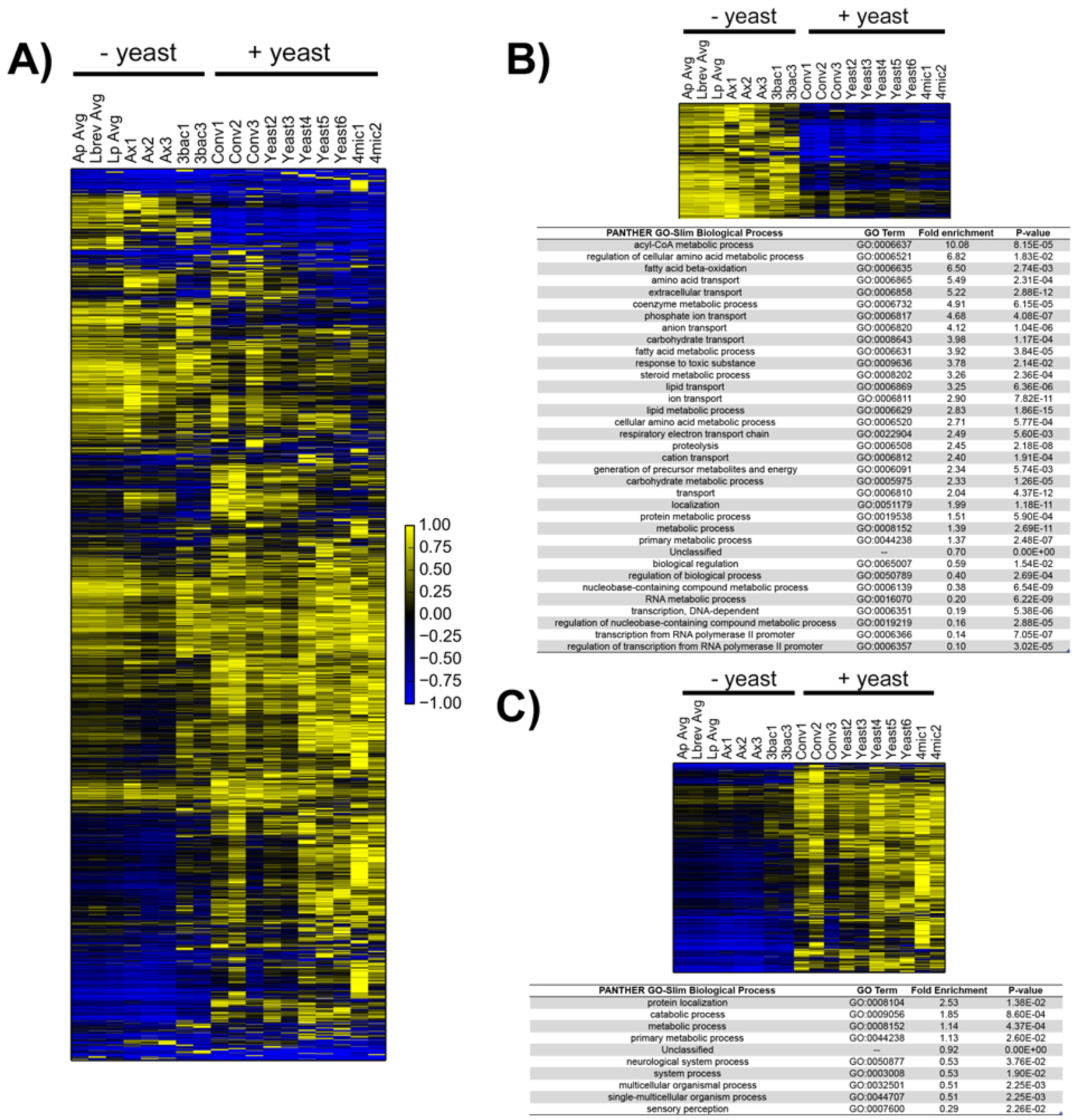
Yeast drive genome-wide difference in gut gene expression. A) Average linkage hierarchical clustering was performed in Gene Cluster 3.0 across all genes that are expressed at least at two FPKM in at least two out of 11 samples. Bacteria mono-association data has been averaged across each treatment to collapse down into a single column. FPKM values for each gene are normalized to range from-1 to 1 before plotting. Abbreviations: Ap avg = average for ***A. pasteurianus***-mono-associated samples; Lbrev avg = average for ***L. brevis***-mono-associated samples, Lp avg = average for ***L plantarum***-mono-associated samples, 3bac = poly-associated (without yeast), Ax = axenic, Conv = conventional, Yeast = ***S. cerevisiae***-mono-associated, 4mic = poly-associated (with yeast). Scale bar is shown at bottom right. B) Top) heatmap of 579 genes that are overexpressed in axenic, bacteria-mono-associated and poly-associated (without yeast) guts compared to other gut samples (Bonferroni p-value>0.05, ANOVA). Bottom) Results from Panther GO-SLIM biological function enrichment test [<u>28</u>] for gene set above compared to reference set of all genes observed across all gut datasets (556 were identified by Panther and used for analysis out of 579) C) Top) Heatmap of 1728 genes that are overexpressed in conventional, yeast mono-associated and poly-associated (with yeast) compared to other gut samples (Bonferroni p-value>0.05, ANOVA). Results from Panther GO-Slim biological processes enrichment test with 1728 (1663 identified) genes compared to reference set of all genes observed across all gut datasets. Note for B) and C): all individual sample values were used for ANOVA analysis, not the average value as plotted in A).

To understand which genes most distinguished these two transcriptional groups, we again performed a gene-by-gene ANOVA. This analysis revealed more than 2,000 genes with significant expression differences between the two groups (p-value under 0.05 after Bonferroni correction). Genes that were more highly expressed in “conventional-like” samples were enriched for annotations for a variety of metabolic processes, most notably involving lipids and amino acids (Figure 3B). Genes more highly expressed in “bacterial-like” samples were also enriched in metabolism annotations, although this enrichment was less pronounced and more general than for the former set (Figure 3C).

Overall, these gut expression data are consistent with a model in which yeast shapes the transcriptional program of the gut in mated, young adult females. Specifically, these results argue that yeast induces metabolic changes that are not evoked by association with any one species of bacteria associated with lab-reared *D. melanogaster* or a simplified community comprised of the all three bacterial species considered herein.

### Global host gene expression in conventional animals is distinct among treatments

As our expectation that host transcription within the gut (i.e. proximal transcription) would change in response to mono-association with different bacterial taxa did not hold true, we were curious if the same would hold true for whole animals. We therefore extracted and sequenced mRNA from individual whole, 5-day post-eclosion, mated female CantonS flies that were prepared with the same microbial treatments as our gut samples (i.e. mono-associated with either *A. pasteurianus, L. brevis, L. plantarum* or *S. cerevisiae*, poly-associated with *A. pasteurianus, L. brevis* and *L. plantarum*, poly-associated with *A. pasteurianus, L. brevis, L. plantarum* and *S. cerevisiae*, axenic or conventional).

From our observation that gene expression in the gut tracked with exposure to yeast, we hypothesized that global transcription would also be dependent on the presence of *S. cerevisiae*. Based on other gene-expression studies examining axenic and conventional animals, we anticipated that the majority of the affected genes still would be those expressed in the gut[<u>11</u>,<u>23</u>].

Data from whole animals (Figure 4A) show that conventionally-reared flies have a transcription profile distinct from all of the samples tested. ANOVA comparing conventional samples to all other samples showed that ^~^1700 genes display different patterns of expression between these two groups after applying a strict Bonferroni correction for multiple testing. Many biological processes are enriched among the genes expressed at higher levels in the conventionally reared samples (Figure 4B), while genes expressed at lower levels show a marked enrichment for processes involving protein folding and biogenesis (Figure 4C).

**Figure 4).**
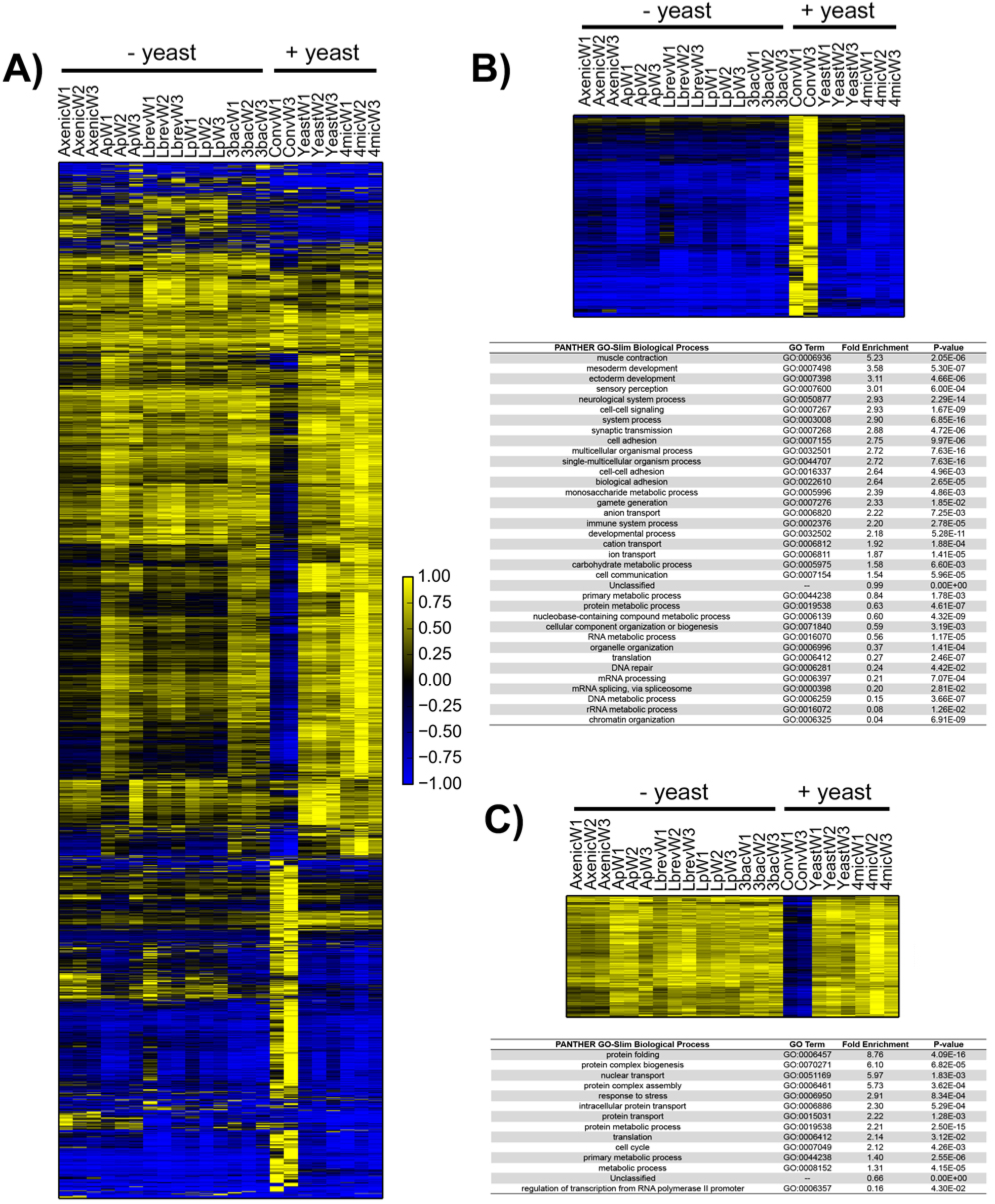
Analysis of gene expression trends from gnotobiotic whole flies (next page) A) Transcriptome-wide heatmap from axenic, conventional, yeast-mono-associated, bacteria-mono-associated and poly-associated whole flies clustered by gene expression Average linkage hierarchical clustering using an uncentered correlation similarity metric was performed in Gene Cluster 3.0 across all genes that are expressed at least at two FPKM across two out of eleven samples. Abbreviations: Ap = ***A. pasteurianus***-mono-associated; Lbrev = ***L. brevis***-mono-associated, Lp = ***L plantarum***-mono-associated, 3bac = poly-associated without yeast, Ax = axenic, Conv = conventional, Yeast = ***S. cerevisiae***-mono-associated, 4mic = poly-associated with yeast. Scale bar is shown at bottom right. B) Top) heatmap of 1159 of 1385 genes that are overexpressed in conventional whole flies compared to other whole fly samples (Bonferroni p-value>0.05, ANOVA). Genes absent in heatmap did not pass filtering criteria. Bottom) Results from Panther GO-SLIM biological function enrichment test [<u>28</u>] for gene set above (1278 genes were identified of 1385) compared to reference set of all genes observed across all whole fly datasets. C) Top) Heatmap 351 that are overexpressed in all non-conventional whole-fly samples compared to conventional whole flies (Bonferroni p-value>0.05, ANOVA). Results from Panther GO-Slim biological processes enrichment test with gene set above (348 genes were identified out of 351) compared to reference set of all genes observed across all whole fly datasets.

Though the global pattern clearly showed that conventional samples exhibit a unique transcription pattern, we reasoned that there could still be a subset of genes that maintain a high similarity between yeast-containing and conventional samples. Gene-by-gene analysis demonstrated that there was only a small overlap in transcription between conventional and yeast-containing samples (^~170^ genes showed similar patterns of expression between samples, (Figure S2) and so better supports a model in which none of the gnotobiotic treatments tested in this study could recapitulate conventional levels of gene expression at the level of the whole animal.

To determine if any differences manifested between different bacterial mono-associated animals at the level of the entire organism that were not apparent in dissected guts, we completed a gene-by-gene analysis as per (Figure 2). As in our guts, there are few significant differences between whole flies that are mono-associated with different bacteria once a Bonferroni correction is applied (Figure S3), though the data do demonstrate more variance than what was observed in dissected guts alone.

Overall, the whole animal data suggest that, while yeast alone may suffice to recapitulate conventional host gene expression in the gut, yeast alone or in a simplified mock community are not enough to generate an animal that demonstrates an overall conventional-like transcriptional program.

### Microbial load is highly variable between lab-reared flies raised under identical gnotobiotic conditions

Prior to conducting the experiments described above, we had been concerned about inter-animal variation in microbial load producing highly variable gene expression patterns. However, the relative lack of variation in gene expression within treatments led us to wonder if our gnotobiotic protocol generated flies with less variability in microbial load than we had anticipated.

We attempted to determine the total bacterial load of the animals we sequenced by counting 16S rRNAs in our sequencing reads. Although our sequencing library preparation protocol involves a polyA selection step to enrich for eukaryotic mRNAs, we nonetheless sequence many *Drosophila* rRNAs, which are not polyadenylated. We therefore expected to have sequenced some microbial rRNAs. However, we did not detect any and, in retrospect, believe this is due to a failure to lyse bacterial cells. Instead, to determine the likely range of microbial loads that our sequenced animals possessed we repeated the preparation of mono-associated animals five times, and determined the microbial loads of multiple animals per treatment per replicate for each of these preparations. Briefly, animals were surface-sterilized, individually homogenized and plated in two dilutions on appropriate media to determine the number of colony forming units (CFUs) they contained (Figure 5A).

**Figure 5).**
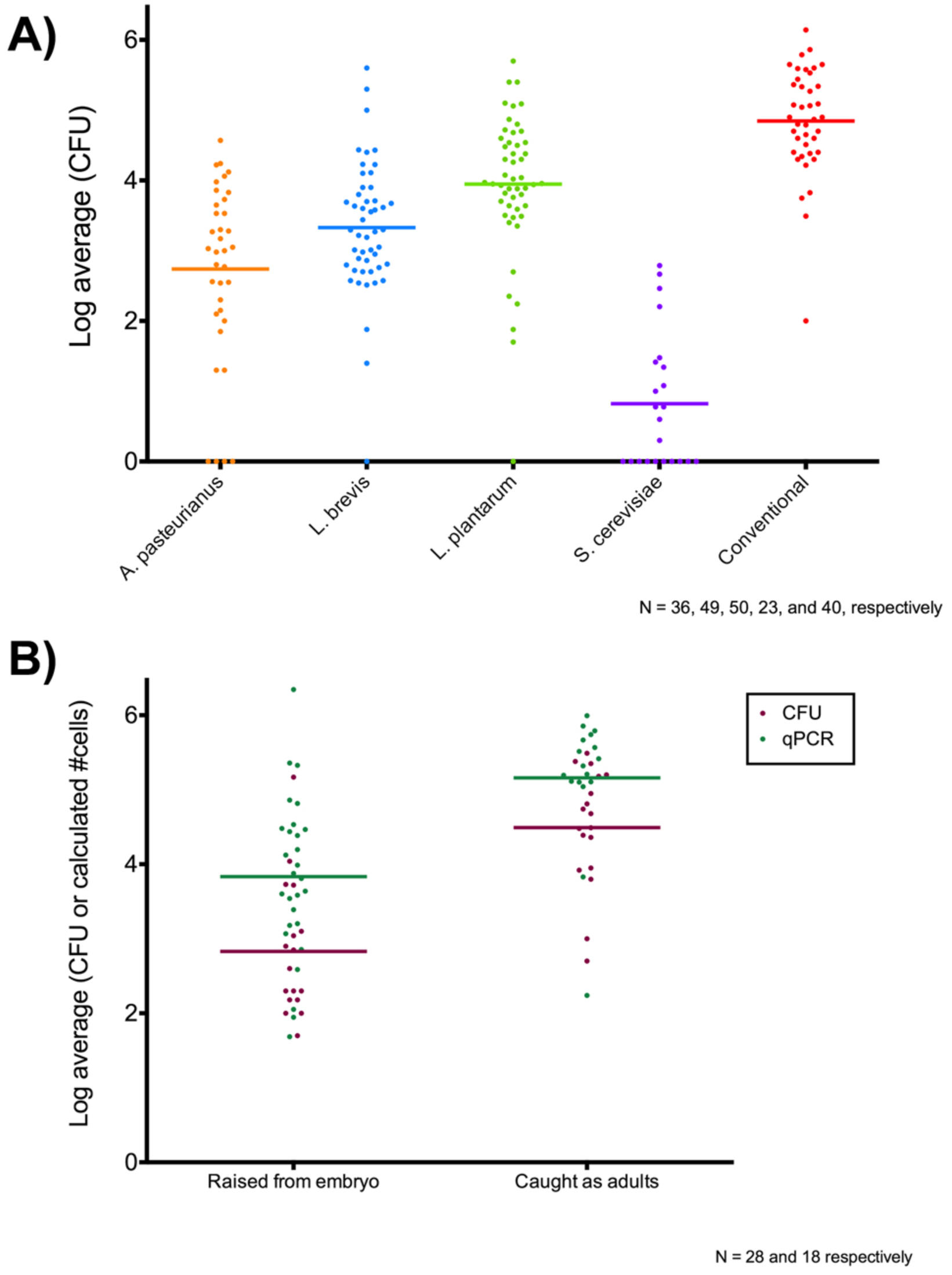
Microbial load of female *D. melanogaster* individuals. A) Log10 transformed average number of colony forming units (CFU) from plating individual gnotobiotic and conventional, lab-reared CantonS, ***Wolbachia***-free, mated, 5-day post-eclosion females on two separate plates. B) Log10 transformed average number of CFU and estimated microbial cells (yeast and bacteria combined) by qPCR for individual, female, wild ***D. melanogaster*** raised from embryos (ranging from three to ten days post-eclosion) or caught as adults (of unknown age). The mean for each group is plotted as a horizontal line.

Consistent with previous work [<u>23</u>], we observed large variability in the microbial load of gnotobiotic flies, despite being reared under nominally identical conditions within a given treatment. The average load among mono-associated animals varies with bacterial species, with *L. plantarum* colonizing more densely than *L. brevis*, which in turn colonizes more densely than *A. pastuerianus*. Conventional animals are on average about 10 - 100x more densely colonized than mono-associated animals. Notably, animals mono-associated with *S. cerevisiae* on average contain only about ten viable yeast cells.

Given these observations we think it is almost certain that the animals we sequenced were effectively colonized with the relevant bacterial species, but that the amount of bacteria they contained was likely highly variable. Despite this presumed variability in the animals we sampled, we did not observe appreciable variation within and between treatments in our transcriptional data. This suggests that variability in microbial load does not play as much of a role in defining transcriptional response as we had anticipated. Still, we cannot completely exclude the possibility that the expression of a few genes did vary in response to microbial load and that we were unable to distinguish this true biological variation from experimental variation.

### Wild-caught flies have comparable microbial loads to lab-reared gnotobiotic flies

We recognized throughout this experiment that our method of preparing gnotobiotic animals is one of many possible methods. Other studies have reported higher inocula for preparing gnotobiotic animals [15,16,29], or inoculating axenic flies upon eclosion rather than associating from embryo [12,19]. We wanted to understand how closely our experimentally-manipulated animals reflected their wild counterparts in terms of gut microbial density. We therefore sampled wild flies from Berkeley, CA and compared their microbial load to that of animals mono-associated in the laboratory. In doing so we had to address several potential complications. It has been well established that microbial load in laboratory *D. melanogaster* positively correlates with age [6,7,16], and thus we wanted to sample animals that would be comparable in age to those used in our sequencing experiments. However, we did not want to capture newly eclosed flies and rear them to the appropriate age in non-natural conditions, as this would deprive them of the substrate needed to replenish their natural gut microbiome and influence their microbial load [30].

We therefore established a stable food source for wild flies using a clean dishpan baited with organic watermelon (referred to as “the fendel”) and then began leaving small pieces of organic fruit (grapes, apple or banana) near, but not touching, the watermelon in the fendel for 24 hours. After 24 hours, the fruit was transferred to a clean vial and any embryos that were deposited were grown up to approximately 5 days PE before surface-sterilization and plating, as done with the gnotobiotic flies. Since our rearing method necessarily involved using a finite number of baits, and we did not know how these baits could impact the final microbial load, we also collected wild flies of unknown age that came to feed in the fendel and subjected them to same plating and DNA extraction protocol. For each fly, DNA was extracted from half of the same homogenate to assay using a non-culture based method (qPCR) to determine microbial load, since some of the taxa within the microbial communities of wild flies might not necessarily grow well under lab conditions, and to confirm species identification, as we wanted only to sample wild *D. melanogaster* (*D. simulans* and *D. immigrans* are morphologically similar to *D. melanogaster*; all three species are found in Berkeley, CA). Culture-dependent and-independent estimations of total microbial loads from flies ranging from 3-10 days and flies of unknown age are shown in Figure 5B.

Our microbial load data showed that five-day old flies are associated with fewer bacteria than wild-caught adults of indeterminate age (presumably older than our young flies and exposed to more substrates as an adult than our reared flies). The qPCR estimates of microbial load was generally higher than our culturing estimates, which is not surprising given the greater diversity of microbial species associated with wild *D. melanogaster* and the expectation that these different species will not necessarily thrive under our chosen culture conditions [2,3]. The number of yeast detected by qPCR was two to three orders of magnitude lower than that for bacteria, consistent with our observation for gnotobiotic animals (Figure S4). As with the gnotobiotic, lab-reared flies, wild fly microbial loads were highly variable even within flies of relatively controlled age. Nonetheless, the association estimates of gnotobiotic and wild flies showed that, with the exception of yeast mono-associated flies, our lab-reared flies bear microbial loads similar to wild flies.

## Discussion

Our results are broadly consistent with previous studies showing that the presence of microbes in the environment and food has a significant effect on gene expression in the adult *D. melanogaster* gut, and we add several new observations to our understanding of the effects of microbes on host gut gene expression. Raising flies in the presence of three bacterial species associated with laboratory stocks of *D. melanogaster* (two of which are the highly abundant in lab-reared flies), either individually or in combination, has minimal effects on adult gut gene expression compared to flies raised axenically. There are, correspondingly, few differences in gut gene expression between flies raised on monocultures of different bacteria. In contrast, the presence of the yeast *S. cerevisiae*, either alone or in combination with bacteria, has a large effect on adult gut gene expression, suggesting that yeast plays a more important role than bacteria in shaping physiology of the adult gut.

### Insensitivity of adult female gut to bacterial species

Contrary to our expectations, we did not find significant differences in gene expression between the guts of *D. melanogaster* individuals mono-associated with three different bacterial species (*A.pasteurianus, L. brevis* or *L. plantarum*) commonly found in lab-reared *D. melanogaster*. This was surprising not only because of previous reports that these mono-associations had phenotypic consequences, but also because we expected, and confirmed, high variability in bacterial load between individuals within the same treatment group. We believe the most parsimonious explanation for our data is that the adult *D. melanogaster* gut is largely indifferent to the identity of bacteria species that occupy it under laboratory colonization levels that reflect those found in wild flies. But several important caveats warrant discussion.

First, that there is no difference in gut gene expression in response to these three strains does not mean that the adult gut is insensitive to bacterial identity. Indeed, while the species found in association with laboratory *D. melanogaster* are also found in wild flies, it is possible that the species that bacterial species that thrive amongst lab-reared flies are not representative of natural populations, and that the *Drosophila* gut would respond in different ways to different bacterial species. We also note that while the *A. pasteurianus* strains used in these experiments was collected from flies, the *Lactobacillus* strains came from non-fly sources (we used the sequenced strains of *L. plantarum* from the ATCC and *L. brevis* from the NBRC), and there may be important inter-strain differences that this choice obscured. Second, although the guts of five day old adult females show no differences between different bacterial mono-associations does not indicate that their guts did not respond differently at earlier points in development; we may simply have missed a critical period for these interactions. In fact, compelling work from several groups has demonstrated that certain fly-associated bacterial strains make significant contributions to larval development [15,16,29] and gut physiology [12,13]. Third, it is possible that the adult female *D. melanogaster* gut is capable of a differential response to these bacteria, but does so only under different conditions (e.g. different diet). Finally, it is possible that the guts in the flies we sampled were responding differently to these bacteria, but that these responses were not reflected at the level of gene expression. For example it could be that the common metabolic demands of the gut dominate its gene expression, and that the gut transduces bacteria-specific signals to other tissues.

Nonetheless, we think it is interesting that gene expression in the adult female *D. melanogaster* gut is so consistently insensitive to varying composition and levels of these bacteria under these conditions. There is a certain logic in insulating the gut from the effects of the constantly varying mix of diverse microbes that wild *D. melanogaster* are exposed to on their preferred substrate of rotting fruits, and our gene expression data may reflect precisely such an insulation. This is not to argue in any way that bacteria in the gut have no effect on the fly, just simply that the epithelial cells of the adult gut do not appear to be the locus for these effects.

### Effect of yeast on gut gene expression

One of the more striking results from this study is the dramatic effect of living *S. cerevisiae* in the environment on gene expression in the adult gut. The strong effect of yeast on *Drosophila* physiology is unsurprising, given the important role that fungi play in the *Drosophila* life cycle, especially as food for larvae. However, the fact that yeast have a large impact on gut gene expression despite the near complete absence of living yeast cells in the gut suggests an indirect effect, either due to differences in development or to the effects of yeast on the nutritional value or other properties of the media. One obvious possible factor is ethanol. We did not perform a chemical analysis of the media, but it is all but certain that the presence of *S. cerevisiae* led to quantifiable ethanol production. *D. melanogaster* preferentially oviposit in substrates emitting volatile fermentation products, and they have evolved a tolerance for relatively high ethanol levels as both larvae and adults. The direct effect of ethanol of gene expression in the *D. melanogaster* gut has not been studied, but dietary ethanol has been shown to have significant effects on gut morphology and physiology [31].

Interestingly, the gut expression data revealed that axenic or bacteria mono-associated guts overexpressed genes involved in fatty acid production and transport relative to yeast mono-associated or conventional guts. This observation is consistent with the finding that axenic animals contain more lipids than conventional animals [10], and suggests that sensation by the gut underlies the amount of fat produced by different gnotobiotic animals.

### Whole animal gene expression paradox

Intriguingly, our experiments showed that, while yeast-containing treatments were sufficient to drive conventional-like transcription in the adjacent host tissue (i.e. guts), yeast alone or as part of a simplified microbial community (*A. pasteurianus, L. brevis, L. plantarum* and *S. cerevisiae*) was not sufficient to produce an animal-wide transcriptional program comparable to that of conventional animals. At first glance, these two findings seem to present a paradox. That is, one might think that if the difference between animals is what microbes are contained in their gut and if the gut is not responding to this change then one should also not observe changes outside of the gut tissue.

We can imagine several ways in which to reconcile these seemingly contradictory results. First, given the repeated observation that different microbial taxa have profound influences on host biology at the larval stages [12,13,15,16,29], we believe that differences in host gut gene expression would likely be observed at earlier timepoints. If this were the case, differences in response to the conventional samples (either due to species, fungal or bacterial, that were not introduced in our simplified community and/or strain-specific effects that could not be recapitulated by our chosen isolates) could lead to differences that accumulate over developmental time and result in animals with markedly different global transcriptional programs. In turn, these accumulated differences could manifest in several ways. The simplest case is that developmental differences accumulated to result in transcriptional differences in an otherwise physically indistinguishable animal. More likely, it could be that certain tissues are over-or underrepresented in conventional animals versus other treatments (which would appear in our data as either higher or lower contributions of those tissues to the overall transcriptional program). Alternatively, the metabolic state of the adult conventional animal could be distinct from those of other treatments (feeding back on transcription to cause big changes).

Lastly, we observed excessive microbial growth on the food in the conventional vials and the conventional animals were much more likely to die between eclosion and collection than any other treatment. It is possible that this sort of stressful environment selected for animals that were acutely adapted to deal with excessive microbial growth, which manifested at the level of global transcription.

### Challenges in utilizing *Drosophila* as a model system to study gut microbe-host biology

Lab-reared flies and their associated bacteria offer a convenient and powerful model for studying microbe-host interactions. However, the diversity and composition of the microbiomes of laboratory flies is limited compared to wild flies [2], likely because we have selected for microbes that thrive on the substrates that we use to culture flies in the lab. A consequence of this selection is that the effects of microbes associated with *D. melanogaster* in the lab may not reflect those of microbes they associate with in the wild.

It is also unclear exactly what it means for microbes to be associated with a fly. Though we generally refer to the microbes found within the fly gut as having colonized the gut, it has yet to be firmly established whether stable colonization of the gut occurs and under what conditions. Analogous to the mucus layer in the mammalian gut, the *Drosophila* gut possesses a barrier, the peritrophic matrix (PM), that impedes the direct contact between objects greater than ^~^250 KDa (including food particles and microbial cells) and host cells [32,33]. Similar to the mammalian mucosal barrier, recent evidence suggests that the PM is a dynamic structure capable of responding to pathogenic microbes [33]. However, while it has been established that some microbes in the vertebrate gut dine on mucus protecting the gut epithelium [34], it has not yet been firmly established whether *Drosophila*-associated microbes actually adhere to or subsist on the molecules comprising or associated with the PM. Additionally, flies appear to lack the crypt structures, present in vertebrate guts, that can provide hideout for microbes allowing them to persist during hard times [35]. At present, we know that lab-reared flies can lose their microbiota by continually transferring onto sterile food [30,36]. The turnover of fly flora under these conditions suggests that stable colonization of the gut tissue is not occurring under these circumstances, although it is possible that there are different conditions under which stable colonization could be observed.

A recent study of wild flies illustrates the importance of microbes to fly biology and highlights the distinct manner in which flies associate with microbial partners. Yamada et al. (2015) discovered that a yeast species, *Issatchenki orientalis*, increases amino acid availability and extends lifespan in flies reared on a protein-deficient diet [36]. They found that lifespan could be extended not just by providing flies with live *I. orientalis*, but also by providing heat-killed *I. orientalis*. Of note, the authors observed that daily transfer of flies onto sterile medium resulted in the loss of association with *I. orientalis*. Thus, there is evidence for flies benefiting from a non-stable association with a microbial species, and the benefit is conferred even when the microbes are dead. This might not be how we typically conceptualize important microbial interactions in animals, but it’s possible that these are the most critical microbial interactions for flies.

## Materials and Methods

### Fly and microbial stocks

*Wolbachia*-free CantonS fly stocks were reared on medium from UC Berkeley’s Koshland fly kitchen (Koshland diet; 0.68% agar, 6.68% cornmeal, 2.7% yeast, 1.6% sucrose, 0.75% sodium tartrate tetrahydrate, 5.6 mM CaCl2, 8.2% molasses, 0.09% tegosept, 0.77% ethanol, 0.46% propionic acid) at 25C. *Acetobacter pasteurianus* (CNE7) was isolated from lab-reared WT *Drosophila melanogaster* and grown on de Man, Rogosa and Sharpe (MRS) agar (Research Products International Corp) at 30C. *Lactobacillus brevis* (NBRC 107147) and *Lactobacillus plantarum* (ATCC 8014) were acquired from NBRC and ATCC culture collections, respectively, and grown on MRS agar at 30C. *Saccharomyces cerevisiae* (ASQ HI) was isolated from wild Hawaiian *Drosophila* by Alli Quan and grown on YPD agar at 30C (A. Quan, personal communication).

### Shotgun DNA sequencing of WT fly guts

Female CantonS flies age five days post-eclosion (PE) were collected via cold anesthesia, surface-sterilized with 10% bleach for ten minutes and rinsed with sterile 1x PBS before dissecting out guts. Forceps used for dissection were treated with 3.5% H2O2 between animals to remove DNA contaminants. Dissections included the proventriculus to the rectal ampulla, leaving the Malpighian tubules attached. Each individual dissected guts was flash frozen in sterile-filtered Buffer ATL (QIAamp Micro Kit, QIAGEN) and stored at −80C. DNA was extracted according to QIAGEN’s QIAamp Micro Kit tissue protocol, with modifications. After the overnight digestion with proteinase K, 0.1 mm Zirconium bead and 1 volume buffer WJL (2M Guanidinium thiocyanate, 0.5 M EDTA, 1.8% Tris base, 8% NaCl, pH 8.5) were added to each sample, Samples were then bead beat twice for one minute at 4C with a 30 second break in between, spun 5 minutes at ^~^14,000xg and the supernatant was transferred to a new tube. Beads were resuspended in two volumes buffer WJL and beat again an additional minute before spinning down and pooling supernatant. Beads were washed once more with two volumes buffer WJL before spinning down and pooling supernatant a final time. An additional spin was performed to pellet any carried-over beads; supernatant was transferred to a new tube. Each sample then received 1 ug of carrier RNA dissolved in buffer AE before proceeding with ethanol precipitation and elution per the manufacturer’s protocol. DNA samples were quantified (Qubit dsDNA HS assay kit, ThermoFisher Scientific) then treated with 120 ng/uL RNase A (QIAGEN) for 1 hour at 37C. Each RNase-treated sample was used to generate an indexed next-generation sequencing library using the TruSeq Nano DNA kit (Illumina). Samples were pooled and sequenced at 100 paired-end reads at the UC Davis Genome Center on a HiSeq 2500. Reads were first filtered by aligning to the *Drosophila melanogaster* genome (version 6.01) using bowtie2, then aligned to the Green Genes 16S database (release 13-5). Species that were identified by two reads or fewer were removed before rarefaction in R (http://www.jennajacobs.org/R/rarefaction.html). Samples that did not rarefy to saturation were discarded.

### Preparation of axenic and gnotobiotic flies

Freshly-laid embryos from CantonS reared on Koshland diet were used to generate axenic, gnotobiotic and conventional animals. Axenic and gnotobiotic animals were prepared using a previously published protocol [10] with some modifications. Briefly, for all treatments, embryos were first collected on molasses plates and rinsed with distilled water. Conventional animals were generated by transferring water-rinsed embryos to YG diet (10% yeast, 10% glucose, 1.2% agar, 0.42% propionic acid, sterilized by autoclaving [37]. To generate axenic and gnotobiotic flies, water-rinsed embryos were subjected to 5 minutes in 10% household bleach, changing bleach solution once halfway through, then briefly rinsed in 70% ethanol before being rinsed with copious distilled water. For axenic animals, cleaned embryos were transferred to sterile YG diet. For gnotobiotic animals, YG vials were inoculated with 50 uL of 1 x 10^8^ cell/mL suspension of one species (mono-associated) or an equal mixture of *A. pasteurianus* (CNE7), *L. brevis* (NBRC 107147) and *L. plantarum* with or without *S. cerevisiae* (HI ASQ) (poly-associated). To prepare cell suspensions, overnight liquid monocultures of microbes were pelleted, washed and resuspended to 1 x 10^8^8^^ cell/mL in 1x PBS following OD600 conversions (given in [10]) for *A. pasteurianus, L. brevis* and *L. plantarum* or 3 x 10^8^ cell/mL per OD600 of 1.0 for *S. cerevisiae* [38].

### Fly collection and validation of gnotobiotic treatment

Gnotobiotic and conventional CantonS females reared on YG diet were collected at age five days PE via cold anesthesia using aseptic technique. For gut samples, individual guts were dissected from each of three female flies (from proventriculus through rectal ampulla including Malpighian tubules) in sterile 1x PBS using Dumont 55 forceps. Cleanly dissected guts were immediately deposited in Trizol (ThermoFisher Scientific) and flash frozen in liquid nitrogen. Between samples, forceps were treated with 70% ethanol, flamed then dipped in 3.5% hydrogen peroxide and sterile water to prevent nucleic acid carryover between samples. For whole fly samples, individual cold-anesthetized animals were transferred to Trizol and frozen as above. *A. pastuerianus, L. brevis* and *L. plantarum* mono-associated animals for gut samples were collected for each of three independent gnotobiotic preparations. All other samples were collected from a single experiment, with preparations for different treatments occurring over the course of several months. Each time animals were prepared and sampled for sequencing three individuals (females when possible, males when not) were independently homogenized and plated on suitable agar medium to verify they possessed (or in the case of axenic animals, lacked) the expected microbial species. For mono-associated animals species verification was achieved by first confirming that plates showed only one morphology, performing 16S or ITS colony PCR on five representative colonies for each treatment and Sanger sequencing of the resultant products. For conventional and poly-associated animals treatment was first verified by checking morphology of resultant colonies performing 16S or ITS colony PCR on five representative colonies for each treatment and Sanger sequencing of the resultant products.

### RNA preparation and sequencing

RNA was prepared from each thawed sample by homogenizing with an RNase-free pestle (Kimble Chase), washing the pestle with 750 uL Trizol, then proceeding using the manufacturer’s protocol with 10 ug glycogen carrier per sample. RNA quality was checked by running on a RNA 6000 Pico chip on a Bioanalyzer 2100 (Agilent Technologies) and quantified using a Qubit fluorometer (Qubit RNA HS assay kit, ThermoFisher Scientific). High quality RNA was then treated with Turbo DNase (ThermoScientific) per the manufacturer’s protocol. For gut samples, RNAseq libraries were prepared using the TruSeq RNA v2 kit (Illumina) starting with 100-200 ng of DNase-treated total RNA for each sample. For whole fly samples, RNaseq libraries were prepared using 400-500 ng of RNA per sample. Samples were multiplexed and sequenced using 100-150 bp paired-end reads on a HiSeq 2500 at the QB3 Vincent J. Coates Genomic Sequencing Facility at UC Berkeley. Reads were aligned to the *D. melanogaster* genome (version 6.01) using Tophat using the “—-no-mixed” option (Table 1). Transcript abundance was calculated from aligned reads using Cufflinks based on a reference transcriptome lacking RNA genes, as we find these to be wildly variable in any RNAseq experiment (M. Eisen, personal communication). Data were analyzed using hierarchical clustering by gene (Cluster 3.0), ANOVA between grouped treatments (scipy.stats) and GO term analysis (Panther, [28]). Data were plotted using matplotlib (Python) and Prism 6 (GraphPad). Heatmap color scales were defined with Chris Slocum’s custom_cmap.py (http://schubert.atmos.colostate.edu/~cslocum/custom_cmap.html).

**Table 1.** Alignment statistics for all mRNAseq libraries reported in the present manuscript.

| Lane | Read type^ | Sample type | Sample ID | Total input read pairs* | Concordantly-aligned pairs* | Concordant alignment rate* |
| --- | --- | --- | --- | --- | --- | --- |
| 1 | 100 PE | Gut | Ap1a | 4.53E+07 | 2.84E+07 | 62.6 |
| 1 | 100 PE | Gut | Ap1b | 2.74E+07 | 1.70E+07 | 61.8 |
| 1 | 100 PE | Gut | Ap1c | 4.96E+07 | 3.10E+07 | 62.5 |
| 1 | 100 PE | Gut | Lbrev1a | 1.39E+07 | 8.61E+06 | 61.9 |
| 1 | 100 PE | Gut | Lbrev1b | 2.08E+07 | 1.30E+07 | 62.2 |
| 1 | 100 PE | Gut | Lbrev1c | 2.06E+07 | 1.22E+07 | 59.1 |
| 1 | 100 PE | Gut | Lp1a | 1.94E+07 | 1.17E+07 | 60.6 |
| 1 | 100 PE | Gut | Lp1b | 2.46E+07 | 1.51E+07 | 61.3 |
| 1 | 100 PE | Gut | Lp1c | 5.41E+07 | 3.53E+07 | 65.2 |
| 2 | 100 PE | Gut | Ap2a | 1.14E+07 | 8.56E+06 | 74.9 |
| 2 | 100 PE | Gut | Ap2b | 1.71E+07 | 1.28E+07 | 75.2 |
| 2 | 100 PE | Gut | Ap2c | 1.84E+07 | 1.39E+07 | 75.7 |
| 2 | 100 PE | Gut | Lbrev2a | 1.34E+07 | 1.05E+07 | 78.0 |
| 2 | 100 PE | Gut | Lbrev2b | 2.37E+07 | 1.80E+07 | 76.0 |
| 2 | 100 PE | Gut | Lbrev2c | 4.64E+07 | 3.55E+07 | 76.5 |
| 2 | 100 PE | Gut | Lp2a | 3.92E+07 | 3.01E+07 | 76.8 |
| 2 | 100 PE | Gut | Lp2b | 2.12E+07 | 1.63E+07 | 76.9 |
| 3 | 100 PE | Gut | Ap3a | 2.19E+07 | 1.70E+07 | 77.4 |
| 3 | 100 PE | Gut | Ap3b | 1.54E+07 | 1.20E+07 | 77.9 |
| 3 | 100 PE | Gut | Ap3c | 2.07E+07 | 1.60E+07 | 77.4 |
| 3 | 100 PE | Gut | Lbrev3a | 2.21E+07 | 1.71E+07 | 77.4 |
| 3 | 100 PE | Gut | Lbrev3b | 1.70E+07 | 1.34E+07 | 78.4 |
| 3 | 100 PE | Gut | Lbrev3c | 2.61E+07 | 2.00E+07 | 76.8 |
| 3 | 100 PE | Gut | Lp2c | 2.05E+07 | 1.57E+07 | 76.5 |
| 3 | 100 PE | Gut | Lp3a | 2.19E+07 | 1.68E+07 | 76.7 |
| 3 | 100 PE | Gut | Lp3b | 3.88E+07 | 3.06E+07 | 78.9 |
| 3 | 100 PE | Gut | Lp3c | 1.57E+07 | 1.22E+07 | 77.5 |
| 4 | 100 PE | Gut | Ax1 | 1.58E+07 | 1.19E+07 | 75.7 |
| 4 | 100 PE | Gut | Ax2 | 3.34E+07 | 2.56E+07 | 76.6 |
| 4 | 100 PE | Gut | Ax3 | 2.96E+07 | 2.28E+07 | 77.2 |
| 4 | 100 PE | Gut | Conv1 | 3.86E+07 | 2.98E+07 | 77.3 |
| 4 | 100 PE | Gut | Conv2 | 2.50E+07 | 1.92E+07 | 77.0 |
| 4 | 100 PE | Gut | Conv3 | 3.33E+07 | 2.54E+07 | 76.4 |
| 4 | 100 PE | Gut | Yeast1 | 3.26E+07 | 2.67E+07 | 81.8 |
| 4 | 100 PE | Gut | Yeast2 | 1.65E+07 | 1.25E+07 | 75.7 |
| 4 | 100 PE | Gut | Yeast3 | 2.10E+07 | 1.59E+07 | 75.5 |
| 5 | 100 PE | Whole | ApW1 | 6.75E+06 | 5.73E+06 | 84.9 |
| 5 | 100 PE | Whole | ApW2 | 7.51E+06 | 6.30E+06 | 83.9 |
| 5 | 100 PE | Whole | ApW3 | 3.77E+06 | 3.23E+06 | 85.6 |
| 5 | 100 PE | Whole | AxenicW1 | 1.05E+07 | 8.91E+06 | 85.0 |
| 5 | 100 PE | Whole | AxenicW2 | 7.64E+06 | 6.45E+06 | 84.4 |
| 5 | 100 PE | Whole | AxenicW3 | 7.97E+06 | 6.71E+06 | 84.3 |
| 5 | 100 PE | Whole | ConvW1 | 2.17E+06 | 1.84E+06 | 84.6 |
| 5 | 100 PE | Whole | ConvW2 | 1.04E+07 | 8.81E+06 | 85.1 |
| 5 | 100 PE | Whole | ConvW3 | 7.14E+06 | 6.06E+06 | 84.9 |
| 5 | 100 PE | Whole | LbrevW1 | 7.77E+06 | 6.69E+06 | 86.1 |
| 5 | 100 PE | Whole | LbrevW2 | 8.81E+06 | 7.60E+06 | 86.3 |
| 5 | 100 PE | Whole | LbrevW3 | 8.03E+06 | 6.88E+06 | 85.7 |
| 5 | 100 PE | Whole | LpW1 | 7.72E+06 | 6.55E+06 | 84.8 |
| 5 | 100 PE | Whole | LpW2 | 1.09E+07 | 9.40E+06 | 86.1 |
| 5 | 100 PE | Whole | LpW3 | 1.22E+07 | 1.05E+07 | 86.4 |
| 5 | 100 PE | Whole | YeastW1 | 1.07E+07 | 9.21E+06 | 86.0 |
| 5 | 100 PE | Whole | YeastW2 | 1.37E+07 | 1.18E+07 | 85.6 |
| 5 | 100 PE | Whole | YeastW3 | 1.37E+07 | 1.16E+07 | 84.8 |
| 6 | 150 PE | Gut | 3bac1 | 4.38E+06 | 2.39E+06 | 54.6 |
| 6 | 150 PE | Gut | 3bac3 | 5.02E+06 | 2.83E+06 | 56.4 |
| 6 | 150 PE | Whole | 3bacW1 | 9.11E+06 | 5.03E+06 | 55.2 |
| 6 | 150 PE | Whole | 3bacW2 | 6.13E+06 | 3.41E+06 | 55.6 |
| 6 | 150 PE | Whole | 3bacW3 | 2.04E+06 | 1.15E+06 | 56.3 |
| 6 | 150 PE | Gut | 4mic1 | 3.53E+06 | 1.97E+06 | 55.8 |
| 6 | 150 PE | Gut | 4mic2 | 4.70E+06 | 2.71E+06 | 57.7 |
| 6 | 150 PE | Gut | 4mic3 | 3.78E+06 | 2.20E+06 | 58.1 |
| 6 | 150 PE | Whole | 4micW1 | 5.86E+06 | 3.21E+06 | 54.7 |
| 6 | 150 PE | Whole | 4micW2 | 6.97E+06 | 3.51E+06 | 50.4 |
| 6 | 150 PE | Whole | 4micW3 | 1.09E+07 | 5.90E+06 | 54.2 |
| 6 | 150 PE | Gut | Yeast4 | 2.80E+06 | 1.60E+06 | 57.0 |
| 6 | 150 PE | Gut | Yeast5 | 3.76E+06 | 2.08E+06 | 55.4 |
| 6 | 150 PE | Gut | Yeast6 | 3.68E+06 | 1.97E+06 | 53.5 |

\*Value obtained from the "alignment\_summary.txt" output following Tophat alignment of reads to v6.01 *D. melanogaster* genome.
^All libraries were sequenced on the Illumina HiSeq 2500 platform

### Determining gut microbe colonization of gnotobiotic flies

Gnotobiotic (mono-associated), axenic and conventional flies were prepared and reared on YG diet five separate times as described above. CantonS females were collected at age five days PE via cold anesthesia using aseptic technique. Animals were surface sterilized using a one minute incubation in 95-100% ethanol then rinsed in sterile water. For conventional and gnotobiotic treatments, a ten flies or as many as available were transferred to individual tubes containing 1x PBS, homogenized with sterile pestles and plated at the dilutions and on the media as shown in Table 2.

**Table 2.** Plating media and dilutions for determining gut microbial load of lab-reared flies.

| Treatment | Plate A | Plate B |
| --- | --- | --- |
| Conventional | 1/200, MRS | 1/200, YPD |
| *A. pasteurianus* | 1/10, MRS | 1/20, MRS |
| *L. brevis* | 1/50, MRS | 1/100, MRS |
| *L. plantarum* | 1/50, MRS | 1/100, MRS |
| *S. cerevisiae* | 1/4, YPD | 1/4, YPD |

For axenic preparations, three females were homogenized in each of two tubes of 1x PBS without prior surface sterilization. One homogenate was plated on MRS and one on YPD. A negative control was performed for each set of flies sampled by homogenizing an equal volume of 1x PBS and plating the half on each MRS and YPD. All plates were incubated at 30C for three days or until plates showed visible colonies. Colonies were counted by hand with the aid of a light board and a handheld counter. The count from each plate was corrected for dilution and averaged over two plates for each fly.

### Determining gut microbe colonization of wild flies

To collect wild flies of unknown age, flies were directly caught either by baiting a closed bottle trap with banana or by directly aspirating from an uncovered plastic dishwashing pan (heretofore referred to as a “fendel”) that was baited with organic watermelon and an assortment of other organic fruits. All baiting and capture was performed in the spring of 2015 at a personal residence in Berkeley, CA. Flies were captured in the morning and transferred into sterile, empty vials (to avoid providing a diet that might manipulate their gut flora) before sampling shortly thereafter. Flies were recovered from vials via cold anesthesia using aseptic technique. Males were discarded.

To collect wild flies of roughly five days PE, a piece of tap-water rinsed organic fruit (grape, banana or fuji apple) was placed in the open-bait fendel and left for 24 hours for provide an oviposition substrate for wild females. After 24 hours, the fruit pieces were placed in sterile vials and the deposited embryos were allowed to develop at room temperature. Vials were monitored daily for newly-eclosed flies. All flies from a vial were collected when the majority of the adults present in the vial were 5 days PE via cold anesthesia using aseptic technique. Males were discarded.

All wild flies were then surface sterilized using a one minute incubation in 95-100% ethanol and rinsed in sterile water before homogenization with a sterile pestle in 1x PBS. Half of the homogenate was diluted to plate 1/200 across MRS agar and YPD agar for each fly and plates were diluted at 30C for three days or until colonies emerged. The remaining half was then processed for DNA extraction as described for shotgun DNA sequencing or frozen at −30C for later processing. A negative control was performed for each set of flies collected by homogenizing an equal volume of 1x PBS with a sterile pestle and splitting in half as for fly samples. Colonies were counted by hand with the aid of a light board and a handheld counter. The count from each plate was corrected for dilution and averaged over two plates for each fly.

All sampled wild females were genotyped by PCR and Sanger sequencing cytochrome c oxidase II primers tLYS (GTTTAAGAGACCAGTACTTG) and tLEU (ATGGCAGATTAGTGCAATGG)[39]. Colonization data from flies that were not identified as *D. melanogaster* (e.g. *D. simulans, D. immigrans, D. persimilis*) were discarded.

To provide a culture-independent method for estimating gut colonization in wild animals, all DNA samples from confirmed wild *D. melanogaster* females were subjected to two sets of triplicate qPCR reactions, one using universal bacteria 16S primer pair P1 (CCTACGGGAGGCAGCAG) and P2 (ATTACCGCGGCTGCTGG) at 100 nM per reaction and the other using fungal ITS primer pair Y1 (GCGGTAATTCCAGCTCCAATAG) and Y2 (GCCACAAGGACTCAAGGTTAG) at 800 nM per reaction [40]. Reactions were performed on a Roche480 LightCycler using Sybr Green mastermix (Roche) templated with 10-20 ng of total fly DNA for each reaction. Four-point standard curves were run for each experiment for *A. pasteurianus* and *L. plantarum* (P1, P2 primer pair) and *S. cerevisiae* (Y1, Y2 primer pair). The amplification program used for qPCR is as follows: 95C for 5 min then 45 cycles of 95 C for 10 seconds, 65C for 10 seconds (decreasing 1 C for the first 10 cycles until reaching 55C), and 72C for 10 seconds. Primers were validated using DNA extracted from monocultures of *A. pasteurianus, L. brevis, L. plantarum* and *S. cerevisiae*. The number of bacterial and yeast cells present in the whole animal was estimated assuming an average genome size of 3 Mb for bacteria and 25 Mb for yeast.

## Supplementary Figures

**Figure S1.**
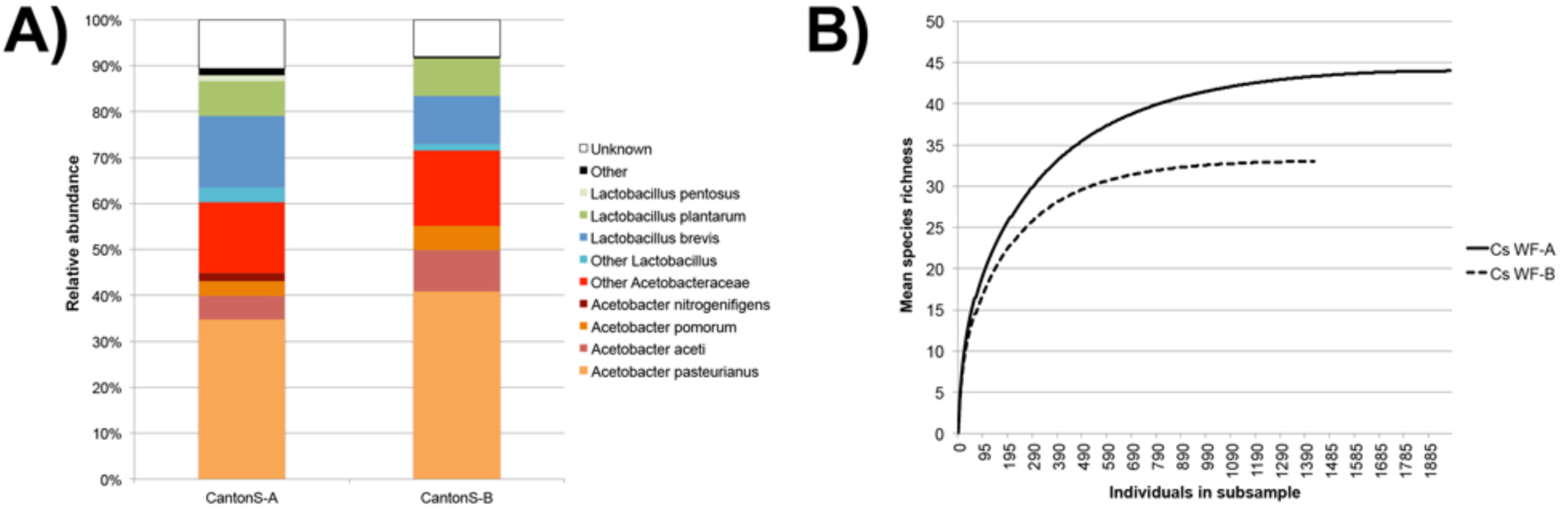
Gut bacterial community analysis of *Wolbachia*-free CantonS five-day mated females reared on Koshland diet. A) Relative species abundance of gut bacteria as determined by shotgun sequencing reads to the Green Genes 16S rRNA database release 13-5. “Other” category includes species not listed in key. “Unknown” category includes reads that aligned to 16S rRNA sequences included in the Green Genes database annotated as “unknown” (e.g. unknown compost). B) Rarefaction curve using data shown in A [0].

**Figure S2.**
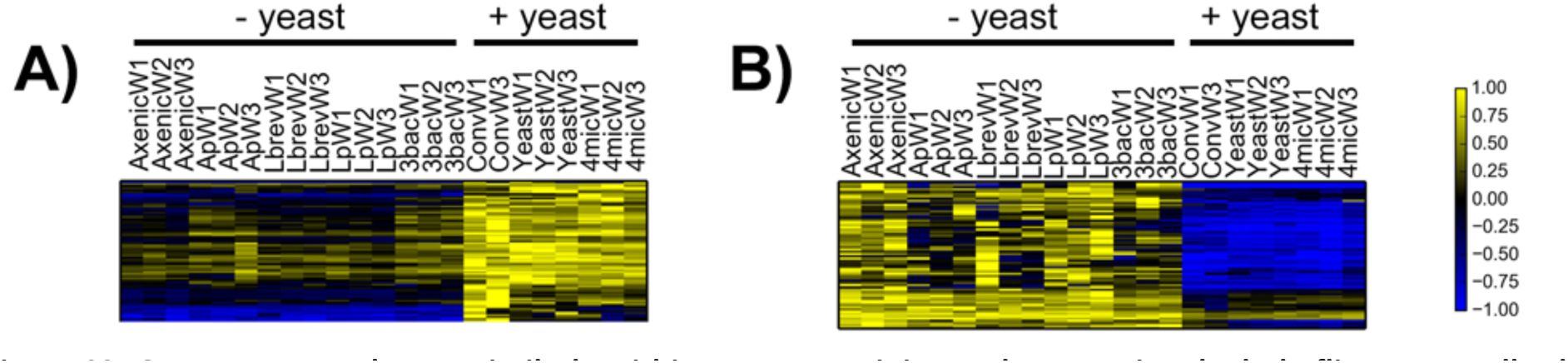
Genes expressed more similarly within yeast-containing and conventional whole flies versus all other samples. **A)** heatmap of 72 genes that are overexpressed in conventional, yeast-mono-and poly-associated whole flies compared to other whole fly samples (Bonferroni p-value>0.05, ANOVA). B) Heatmap of 67 genes that are overexpressed in axenic and bacteria mono-and poly-associated whole flies compare to other whole fly samples (Bonferroni p-value>0.05, ANOVA). Abbreviations: Ap = ***A. pasteurianus***-mono-associated; Lbrev = ***L brevis***-mono-associated, Lp = ***L plantarum***-mono-associated, 3bac = poly-associated without yeast, Ax = axenic, Conv = conventional, Yeast = ***S. cerevisiae***-mono-associated, 4mic = poly-associated with yeast.

**Figure S3.**
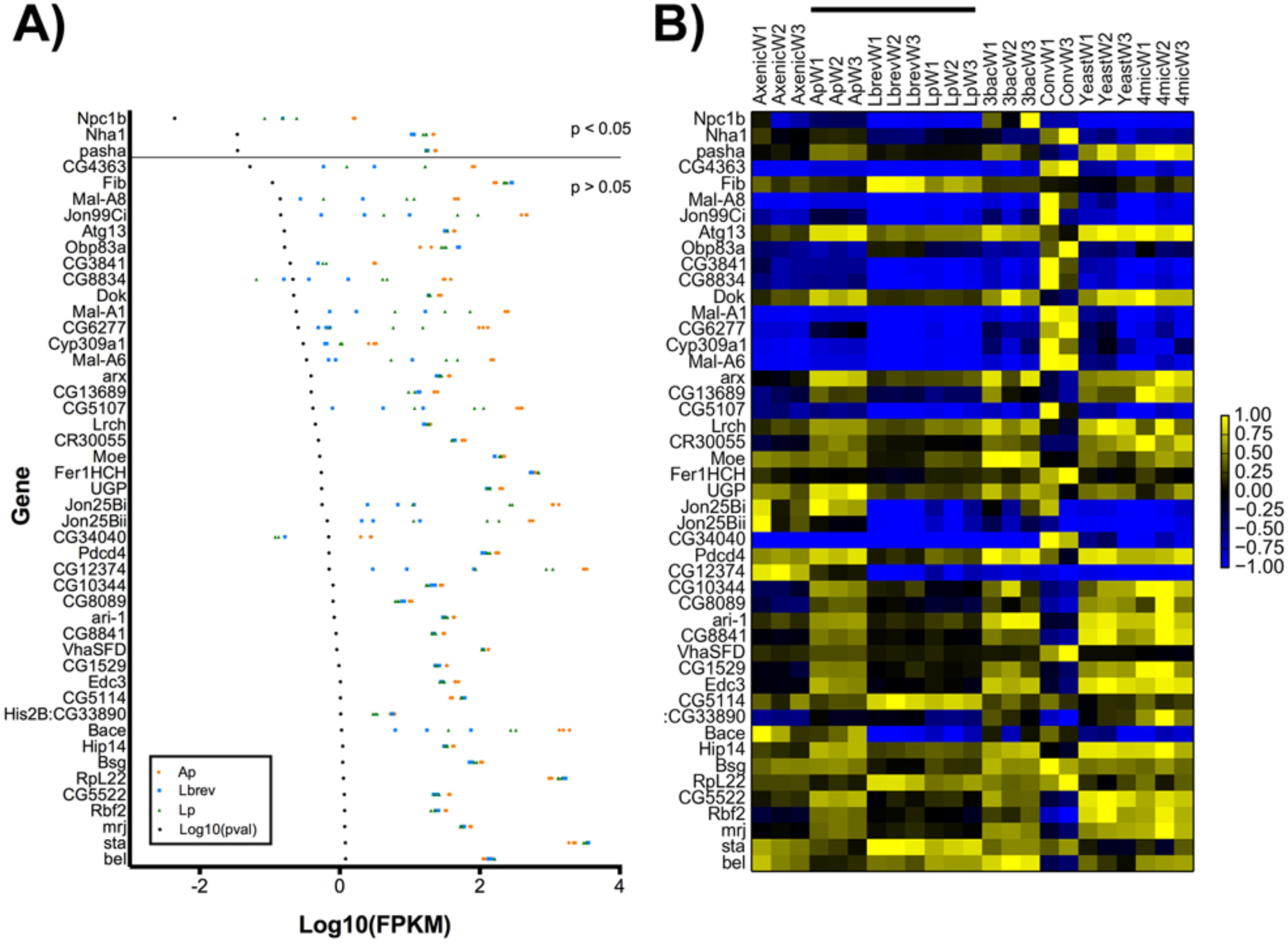
Genes showing greatest difference in expression values between different bacteria mono-associations in whole adults as determined by one-way ANOVA. A) Scatterplot of log10-transformed FPKM values for each bacteria mono-associated whole fly replicate. Genes are ordered from lowest ANOVA p-value (top) to highest (bottom). P-values have undergone a Bonferroni correction for multiple testing. B) Data from A presented as a heatmap. FPKM values for each gene are normalized to range from-1 to 1 before plotting. Black line above heatmap denotes bacteria mono-association samples. Abbreviations: Ap = ***A. pasteurianus***-mono-associated; Lbrev = ***L. brevis***-mono-associated, Lp = ***L plantarum***-mono-associated, 3bac = poly-associated without yeast, Ax = axenic, Conv = conventional, Yeast = ***S. cerevisiae***-mono-associated, 4mic = poly-associated with yeast.

**Figure S4.**
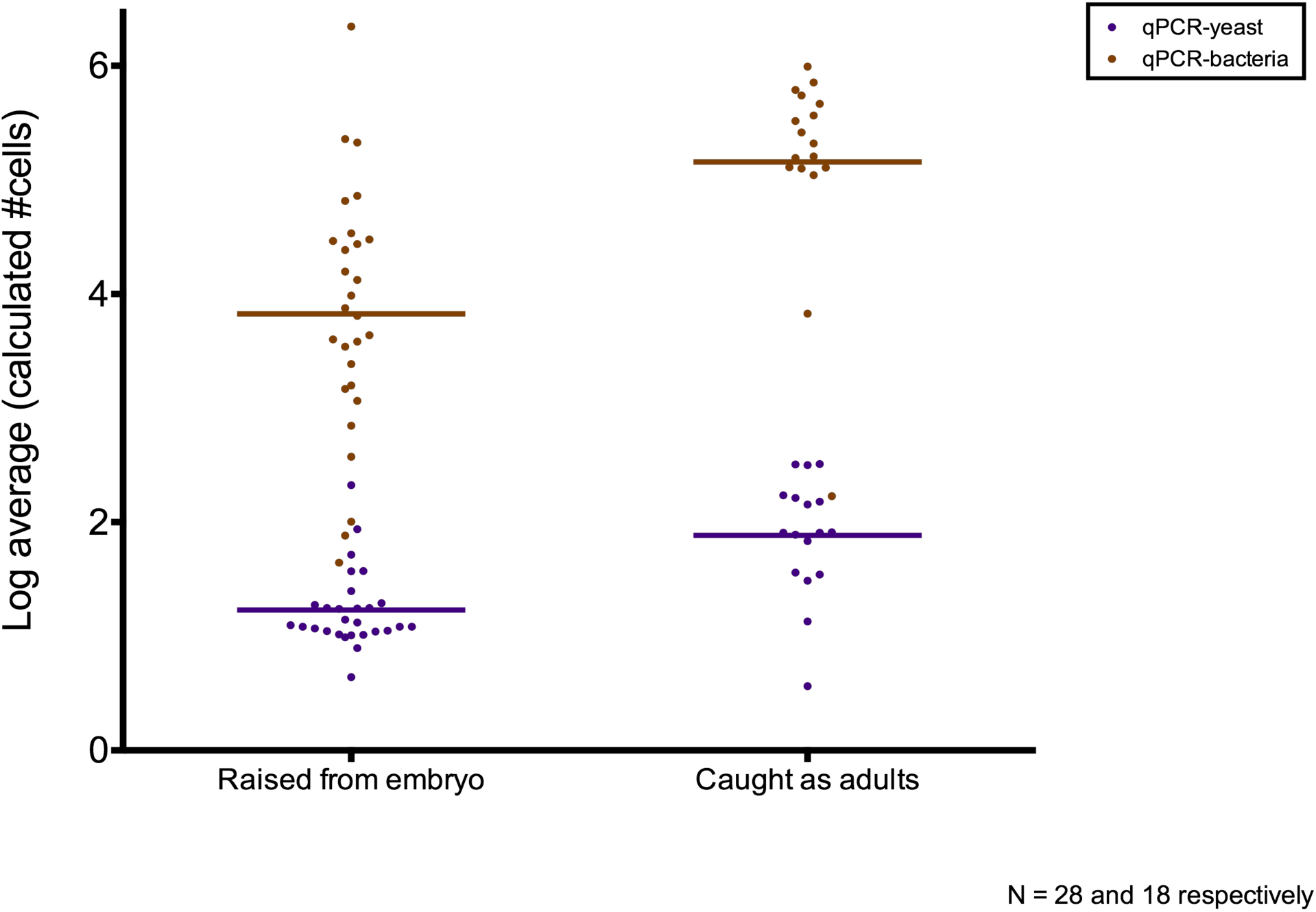
Loads of bacteria and yeast in wild flies estimated from qPCR. Log10 transformed average number of estimated bacteria cells or yeast cells by qPCR for individual, female, wild ***D. melanogaster*** raised from embryos (ranging from 3-10 days post-eclosion) or caught as adults (of unknown age). The mean for each group is plotted as a horizontal line.

